# Malaria in Pregnancy and Endotoxemia Disturb Maternal Thyroid Function and Remodel the Expression of Placental Thyroid Hormone Transporters in Mice

**DOI:** 10.64898/2026.11.24.768855

**Authors:** Klaus Novaes Fontes, Cherley Borba Vieira de Andrade, Verônica Müller de Oliveira Nascimento, Victoria Regina de Siqueira Monteiro, Hanailly Ribeiro Gomes, Alessandra Cristina Chagas Valim, Douglas Esteves Teixeira, Ana Acacia Sá Pinheiro, Flavia Fonseca Bloise, Enrrico Bloise, Tania Maria Ortiga-Carvalho

## Abstract

Thyroid hormones (TH) are essential for fetal development. We hypothesized that malaria in pregnancy (MiP) and lipopolysaccharide (LPS) challenge would disrupt the expression of placental TH transporters. Pregnant dams were infected with *Plasmodium berghei ANKA* at gestational day (GD) 13.5 or challenged with lipopolysaccharide (LPS) at GD17.5. Maternal thyroid parameters, fetal outcomes, placental inflammatory markers, TH transporters and deiodinases expression were assessed at GD18.5. Both MiP and LPS increased maternal serum thyroid-stimulating hormone (TSH) and reduced fetal weight, whereas MiP decreased free thyroxine (fT4). MiP reduced fetal size and the fetal-to-placental weight ratio. Placental *Cxcl1* mRNA increased in both sexes following MiP and in females following LPS injection, whereas *Ccl2* mRNA increased only in males following LPS challenge. MiP reduced *Slc7a8* mRNA in both sexes and *Dio2* mRNA in males. No other significant changes in the assessed placental transcripts were detected in either model. At the protein level, MiP increased MCT8 immunostaining in the labyrinth zone (Lz) in both sexes and in the junctional zone (Jz) in males. MCT10 immunostaining decreased in the Lz and increased in the Jz in both sexes following both challenges. MiP increased LAT2 and OATP4A1 immunostaining in both zones in males. Following LPS challenge, MCT8 immunostaining increased in both zones in males, LAT2 increased in the Lz in males, and OATP4A1 increased in the Lz in both sexes. In conclusion, MiP and LPS-challenge disturb maternal thyroid and reshape placental TH transporter expression in a sex- and zone-dependent manner.

**Subject Area:** Endocrinology and Metabology

**NEW FINDINGS:** *What is the central question of this study?:* How do Malaria in Pregnancy (MiP) and acute endotoxemia affect placental TH metabolism?

*What is the main finding and its importance?:* MiP and acute endotoxemia induced by LPS generate distinct disturbances in the expression of placental TH transporters, local inflammation and maternal thyroid dysfunction, which may, to some extent, affect the placental regulation of iodothyronine transport and metabolism and contribute to adverse gestational outcomes such as fetal growth restriction and preterm delivery.

## INTRODUCTION

Gestational infections induce marked alterations in the physiology of the maternal-fetal interface, resulting in adverse outcomes such as miscarriage, intrauterine growth restriction (IUGR) and preterm delivery (PTD) (Derricott *et al*., 2013; Chawanpaiboon *et al*., 2019). Malaria in pregnancy (MiP) and intrauterine dysbiosis related to bacterial infections deserve special attention, since both conditions trigger the synthesis of inflammatory mediators in intrauterine tissues. These mediators can potentially impair the expression of key placental transporter systems, leading to the development of IUGR and associated neonatal comorbidities (Challis *et al*., 2009; Boeuf *et al*., 2013).

Globally, approximately 125 million pregnancies per year occur in malaria-endemic areas and are therefore at risk of MiP. The main pathophysiological characteristic of MiP is the placental sequestration of *Plasmodium*-infected erythrocytes, an event that triggers chronic systemic and local inflammatory responses impairing the transport of nutrients between the mother and the fetus (Neres *et al*., 2008; Conroy *et al*., 2013). Therefore, MiP is associated with an increased risk of maternal/fetal mortality, IUGR, low birth weight, miscarriage, and PTD (Rogerson *et al*., 2018).

Ascending vaginal infection is an important pathway in which bacterial species from the lower genital tract gain access through the cervix into the uterine cavity, leading to dysbiotic colonization of the placenta and fetal membranes (Suff *et al*., 2018) and increasing the risk of PTD and IUGR(Goldenberg *et al*., 2008; Li *et al*., 2012). Such infections are frequently associated with dysbiotic colonization of amniotic fluid (Romero *et al*., 2019), chorioamnionitis, and postpartum endometritis (Goldenberg *et al*., 2008). Additionally, bacterial infections that affect the maternal-fetal interface can also be related to the hematogenous route, commonly arising from maternal periodontal disease and urinary tract infections (Goldenberg *et al*., 2008).

The placenta expresses thyroid hormone (TH) transporters, including the monocarboxylate transporters MCT8 and MCT10, the organic anion transporting polypeptide, OATP4A1, and the L-type amino acid transporters LAT1 and LAT2, in both the apical and basolateral membranes of the human placental syncytiotrophoblast, with corresponding expression in the murine labyrinthine zone (Lz) (James *et al*., 2007; Chen *et al*., 2022). The expression of the iodothyronine deiodinases is also described, with D3 being predominantly localized at the apical membrane of the syncytiotrophoblast and D2 at the basolateral membrane of cytotrophoblast cells (Chan *et al*., 2009). Changes in the transport and metabolism of TH can also be directly related to consequences for the developing fetus, especially to IUGR and neurodevelopmental sequelae (Loubière *et al*., 2010; Azhan *et al*., 2026).

TH regulate numerous physiological processes related to proper placental development and function, including those related to early placentation (Wu *et al*., 2019) and the invasive potential of extravillous trophoblasts (Oki *et al*., 2004; Appiah Adu-Gyamfi *et al*., 2020). TH also induce the expression of angiogenic and inflammatory factors involved in spiral arteries remodelling (Appiah Adu-Gyamfi *et al*., 2020). Finally, recent evidence suggests the participation of iodothyronines, primarily through the non-genomic action pathway, in the activation of placental inflammasomes (Chen *et al*., 2015; De Luca *et al*., 2021), which, when dysregulated, are directly involved in the pathogenesis of preeclampsia and IUGR (Menkhorst *et al*., 2023). Even though it is well established how important TH are for the maintenance of a successful pregnancy and fetal central nervous system development, there is still limited information about how *Plasmodium berghei* ANKA (modelling MiP) or bacterial lipopolysaccharide (LPS; modelling bacterial infection) impact placental TH transporter systems and metabolism.

In the present study, we used previously established and validated murine models of MiP and LPS-induced bacterial endotoxemia (Bloise *et al*., 2013; Fontes *et al*., 2019; Martinelli *et al*., 2020a, 2020b; Reginatto *et al*., 2021) to investigate how *Plasmodium berghei ANKA* infection and LPS exposure affect maternal TH status, placental TH transporters and deiodinases expression, and the local placental inflammatory response in pregnant C57BL/6 dams. Given the essential roles of TH in placental function and fetal growth/neurodevelopment, we hypothesized that these two inflammatory challenges would differentially alter maternal TH status and the expression of key placental TH transporters and deiodinases at term, with effects varying according to fetal sex and placental compartiment.

## MATERIALS AND METHODS

### Animal Treatment and Experimental Design

Female and male C57BL/6 mice were kept under a temperature-controlled environment (23°C), with a 12-hour light/dark cycle, with *ad libitum* access to food and water. A total of 48 virgin females (8 to 10 weeks of age) underwent a hormonal stimulation protocol for ovulation (Lamas *et al*., 2021) and were bred with males (10 to 12 weeks of age) at a ratio of 2 to 3 females for each male, as previously described (Bloise *et al*., 2013; Fontes *et al*., 2019; Reginatto *et al*., 2021; Monteiro *et al*., 2022). Detection of vaginal plugs the morning after mating was marked as gestational day 0.5 (GD 0.5). A body weight gain of 3 g to 5 g between GD0.5 and GD13.5 was used to confirm pregnancy in both models. The criteria used for animal randomization, following pregnancy confirmation, included assigning the animals to experimental groups based on the number of pregnant females and their respective body weights. The sample size in each experiment was determined based on previous studies conducted by our group (Fontes *et al*., 2019; Reginatto *et al*., 2021).

The present study was approved by the Animal Care Committee of the Health Sciences Center, Federal University of Rio de Janeiro (CEUA protocol number: 128/22) and registered with the Brazilian National Council for Animal Experimentation Control. Humane laboratory animal care followed the “Principles of Laboratory Animal Care” proposed by the National Society for Medical Research and the U.S. National Academy of Sciences Guide for the Care and Use of Laboratory Animals.

Our MiP murine model was performed as previously described (Fontes *et al*., 2019; Martinelli *et al*., 2020a). Briefly, at GD13.5, an intraperitoneal (i.p.) injection of *Plasmodium berghei* ANKA-infected erythrocytes (5×10^5^ infected erythrocytes, n=13; obtained from animals exposed to cryopreserved protozoa), or phosphate buffered saline (PBS-control; n=10) was administered to pregnant C57BL/6 mice. Animals were euthanized at GD18.5 and maternal blood, fetuses, and placentas were collected.

Our endotoxemia model was conducted as described previously (Martinelli *et al*., 2020b; Reginatto *et al*., 2021). Briefly, an intraperitoneal (i.p.) injection of LPS (Sigma, *E. coli* O55:B5, 150 µg/kg) or PBS (control group) was administered to pregnant mice at GD17.5. Animals were euthanized at GD18.5, 24h (n=10 for control group and n=15 for LPS group) after the injection of a single dose of LPS/PBS and maternal blood, fetuses and placentas were collected.

At GD18.5, before euthanasia, blood from the caudal vein was collected and Giemsa-stained thin blood smears were performed to determine maternal parasitemia. Then, using light microscopy, the number of infected erythrocytes per erythrocytes counted in 10 fields was determined. Animals were anesthetized, on GD18.5, with isoflurane (Cristalia, São Paulo, Brazil) and euthanized by decapitation. Cardiac puncture was performed to collect maternal blood, which was centrifuged (1,077 x g, 15 min) and stored at −80°C until the analysis for fT4 and TSH serum levels.

Maternal spleen, placental discs, and fetuses were dissected and weighed on an analytical balance. Two placentas per litter, one for each fetal sex (determined by external genitalia morphology) (Murdaugh *et al*., 2018) with wet weights nearest to the litter mean(Bloise *et al*., 2012), were selected and cut in half, using the umbilical cord insertion as reference (Bloise *et al*., 2012; Connor *et al*., 2020). One half was stored at −80°C for further qPCR analysis, while the other half was fixed overnight in 4% paraformaldehyde (Sigma-Aldrich, Missouri, USA) for protein expression and localization via immunohistochemistry. Figure 1 shows the charts of experimental designs for both gestational infection models used in the present study.

**Figure 1.**
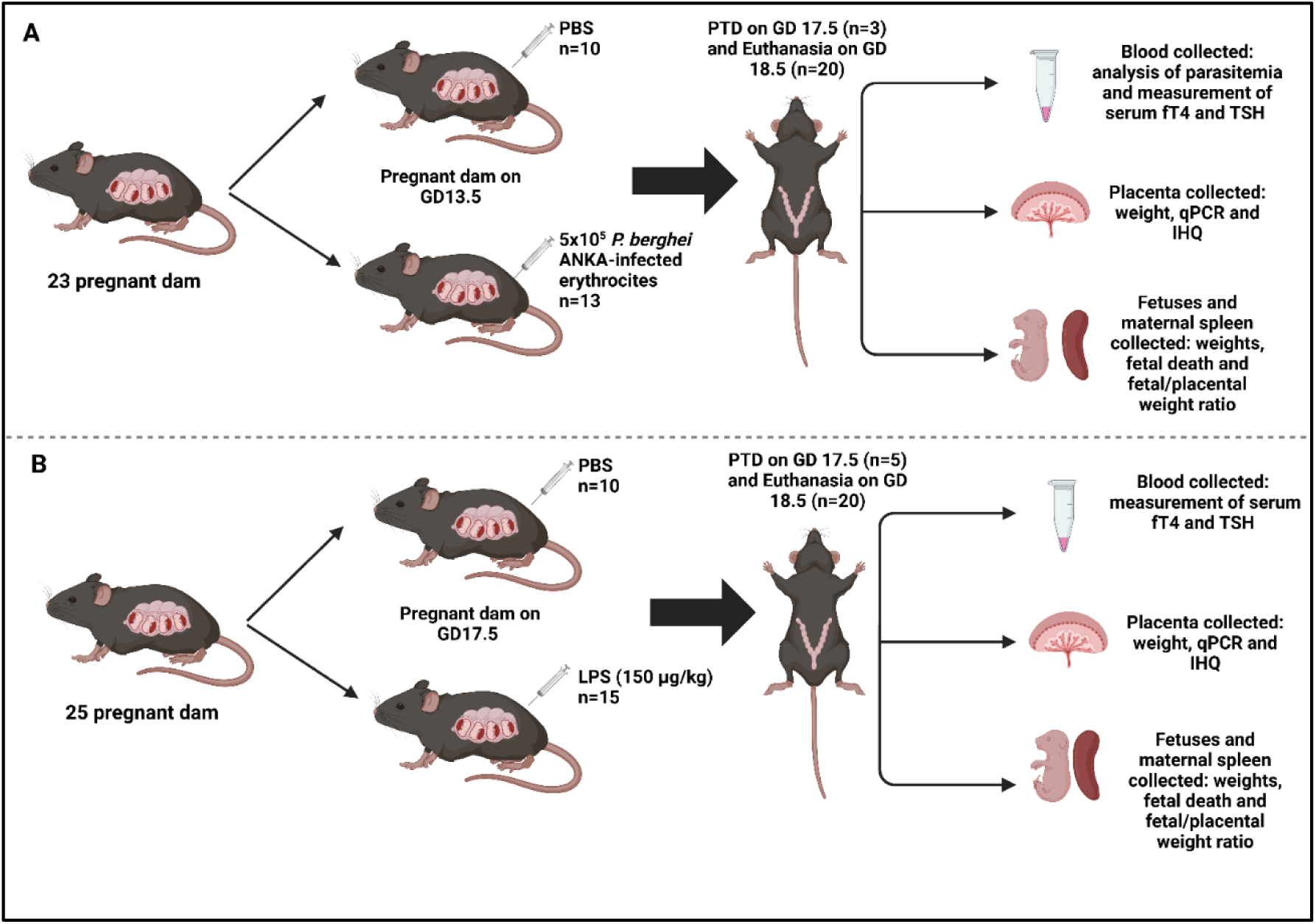
*Plasmodium berghei* ANKA infection and LPS administration during pregnancy in C57BL/6 mice. (A) Chart of the experimental design in MiP experiment. **(B)** Chart of the experimental design in LPS model. Created in BioRender. Fontes, K. (2026) https://BioRender.com/zoooq0y

### qPCR

Total RNA was isolated from one half of each placental disc using the TRIzol reagent (TRIzol reagent; Life Technologies, California, USA) following manufacturer recommendations. RNA concentration and purity were assessed using the NanoDrop One Spectrophotometer (Thermo Fisher Scientific, Massachusetts, USA), in which absorbance values were obtained at 260 nm, in addition to considering the A260/A280 ratios ranging between 1.8 and 2.0. RNA integrity was confirmed through gel electrophoresis. cDNA was synthesized from total RNA (1 µg) using the High-Capacity cDNA Reverse Transcription Kit (Applied Biosystems, São Paulo, Brazil). The mRNA levels of selected TH transporters, deiodinases and pro-inflammatory chemokines (see Table 3 for the primer sequences) were quantified by qPCR using the Maxima SYBR Green/Rox qPCR Master Mix 2x and the QuantStudio 3 Real-Time PCR System (Thermo Fisher Scientific, Massachusetts, USA). Cycling conditions combined an initial step at 50°C (1 step of 2 min) and an initial denaturation at 95°C for 10 min, followed by 40 cycles of denaturation at 95°C (15 s), primer annealing at 60°C (30 s), and extension at 72°C (45 s). The expression of target genes was normalized to the geometric mean of the following reference genes: *Gapdh, Ywhaz and Ppib*, which expression was stable across groups in both models (Table 3). Relative expression was calculated using the 2^-ΔΔCt^ method (Livak & Schmittgen, 2001). All samples and standards were measured in duplicate and only assays with efficiencies ranging between 95 and 105% were considered acceptable for analysis. Contamination by exogenous genomic DNA was avoided by using intron-spanning primers, reverse transcriptase-negative samples and melting curve analyses obtained from each qPCR reaction.

### Histological tissue processing and Immunohistochemistry

Twenty fixed placentas (4% paraformaldehyde) of each experimental group (five placentas from both male and female offspring, either in the control or malaria/LPS groups) underwent a histological processing protocol with increasing concentrations of ethanol (Isofar, Rio de Janeiro, Brazil), diaphanization in xylol (Isofar, Rio de Janeiro, Brazil) and inclusion in paraffin (Easypath, São Paulo, Brazil). Two Tissue Microarray TMA blocks were generated (Jawhar NMT, 2009): one containing only labyrinth zone (Lz) tissues and the other containing only junctional zone (Jz) tissue. Each TMA slide contained forty placental sections (5µm thickness, twenty sections from each experimental model) obtained using a Rotatory Microtome CUT 5062 (Slee Medical GmbH, Nieder-Olm, Germany), which were subsequently subjected to immunohistochemistry.

Following deparaffinization and rehydration in graded ethyl alcohol (100%, 90% and 70%), sections were incubated in 3% hydrogen peroxide (endogenous peroxidase blocking). PBS+Tween was used to remove excess hydrogen peroxide. The antigen retrieval was performed by immersing the slides in Tris-EDTA buffer (pH 9.0, 15 minutes in the microwave), followed by immersion in sodium citrate buffer (pH 6.0, 8 minutes in the microwave). The blocking of non-specific antibody binding sites was achieved by incubating the slides in bovine serum albumin (3%) in PBS for 1 hour. Then, the slides were incubated overnight at 4°C with the following primary antibodies: MCT8 (ab192828, 1:200; Abcam Plc, Cambridge, UK), MCT10 (PA5-48538, 1:50; Thermo Fisher Scientific, Massachusetts, USA), LAT2 (PA5-62519, 1:200; Thermo Fisher Scientific, Massachusetts, USA) and OATP4A1 (PA5-56535, 1:200; Thermo Fisher Scientific, Massachusetts, USA). The next day, the slides were incubated for 1 hour with a pan-specific universal biotin conjugated secondary antibody (VectorLabs, United States). Followed by three washes with PBS (pH 7.2) for 5 minutes and last, incubated with peroxidase-streptavidin complex (VectorLabs, United States) for 30 minutes. The stained slides were developed with 3,3-diamino-benzidine (DAB; VectorLabs, United States). After immunostaining, tissue sections were counterstained with hematoxylin (1 minute), dehydrated in increasing concentrations of ethanol (70%, 90% and 100%), immersed in xylol (three xylene immersions) and mounted with cover slips and Entellan (Merck Millipore, Massachusetts, USA). Negative controls for immunohistochemistry were prepared by omitting primary antibodies. The slices were scanned using the NanoZoomer S360 (Hamamatsu Photonics K. K, Japan) and the images were obtained using the NDP.view2 software. Quantification of MCT8, MCT10, LAT2, OATP4A1 was performed using the mask tool of the Image Pro Plus 5.0 software (Media Cybernetics, Maryland, USA), considering only the percentage of positively stained cells relative to the total viable tissue area (empty spaces of the images were excluded) (Fontes *et al*., 2019; Andrade *et al*., 2021, 2026; Reginatto *et al*., 2021). Two investigators, blinded to the experimental groups and models, analyzed a total of fifteen random field digital images captured per tissue fragment of each placental zone (labyrinth and junctional zone) in 10 control, 10 malaria and 10 LPS-injected mice (5 male and 5 female mice, for both experimental groups), totaling 1200 digital images analyzed for each TH transporter. The average of the fifteen fields was considered the single representative value for a specific placental area of each animal (Andrade *et al*., 2021, 2026).

### Measurement of serum fT4 and TSH levels

Maternal serum free thyroxine (fT4) and thyroid stimulating hormone (TSH) concentrations were measured using the following commercially available ELISA kits: Mouse TSH (Thyroid Stimulating Hormone) ELISA Kit (EM1433; Fine Test, Wuhan, China) and Mouse fT4 (Free Thyroxine) ELISA Kit (EM1037; Fine Test, Wuhan, China), according to manufacturer’s datasheet recommendations. The measurement range for each test was 1.25-80 ng/ml for TSH and 1.563-100 pg/ml for fT4. Intra and inter-assay variations were 4.8% and 5.9%, respectively, for TSH, and 4.7% and 4.9%, respectively, for fT4. Serum analyses for each analyte, in both experimental models, were performed on the same day, by the same operator, and using the same plate. The optical densities in both assays were measured by reading the absorbance at 450 nm in the Emax Plus Microplate Reader (Molecular Devices, California, USA). The mean absorbance values for each standard, control and sample were calculated and plotted on a logistic standard curve, allowing the calculation of fT4 and TSH concentrations in each sample.

### Statistical analysis

All data are expressed as the mean ± standard error of the mean (SEM). Normal distribution was evaluated by D’Agostino-Pearson tests. The Student’s T-test or non-parametric Mann-Whitney test were, then, applied according to the results of normality tests. The statistical parameters of pregnancy were obtained by using the mean value of all fetuses in a litter for each C57BL/6 dam, and not the individual fetuses. Consequently, “n” represents the number of litters (Festing, 2006; Coan *et al*., 2008; Bloise *et al*., 2012). Proportions, namely preterm delivery and fetal death rates were copared using Fisher’s exact test. Dams that underwent preterm delivery between GD17.5 and GD18.5 were excluded from the analyses of litter size, fetal and placental parameters, and gene and protein expression. For qPCR and immunohistochemistry data, closest placentas (from both male and female fetuses) to the mean weight of all placentas were selected from each litter. Statistical analyses were performed using the GraphPad Prism 8 software (GraphPad Software, California, USA), and differences were significant at *P*<0.05.

## RESULTS

### MiP causes maternal morbidity and fetal growth restriction, whereas LPS selectively reduces fetal weight

In the MiP experimental model, the average parasitemia in *Plasmodium*-infected dams at GD18.5 was 15.7% (range 5-30%). Pregnant malaria-infected mice had reduced body weight gain between GD13.5 and GD18.5 (*P*=0.003) and increased spleen weight (*P*<0.001), compared to the control group. Fetal weight (*P*<0.001), fetal size (*P*=0.020), and fetal:placental (F:P) weight ratio (*P*=0.012), a well-established index of placental efficiency (Wilson & Ford, 2001), were reduced in malaria-infected dams (Table 1). No changes were observed in the litter size (*P*=0.071) and in the placental weight (*P*=0.200) of malaria-infected dams, compared to control group. Furthermore, our murine model of MiP promoted a PTD rate of 23.1% (3 of 13 infected dams had vaginally born dead fetuses between GD17.5 and GD18.5) and a fetal death rate of 19.6% (10 out of 51 fetuses of infected dams showing clear signs of non-viability).

**Table 1:**
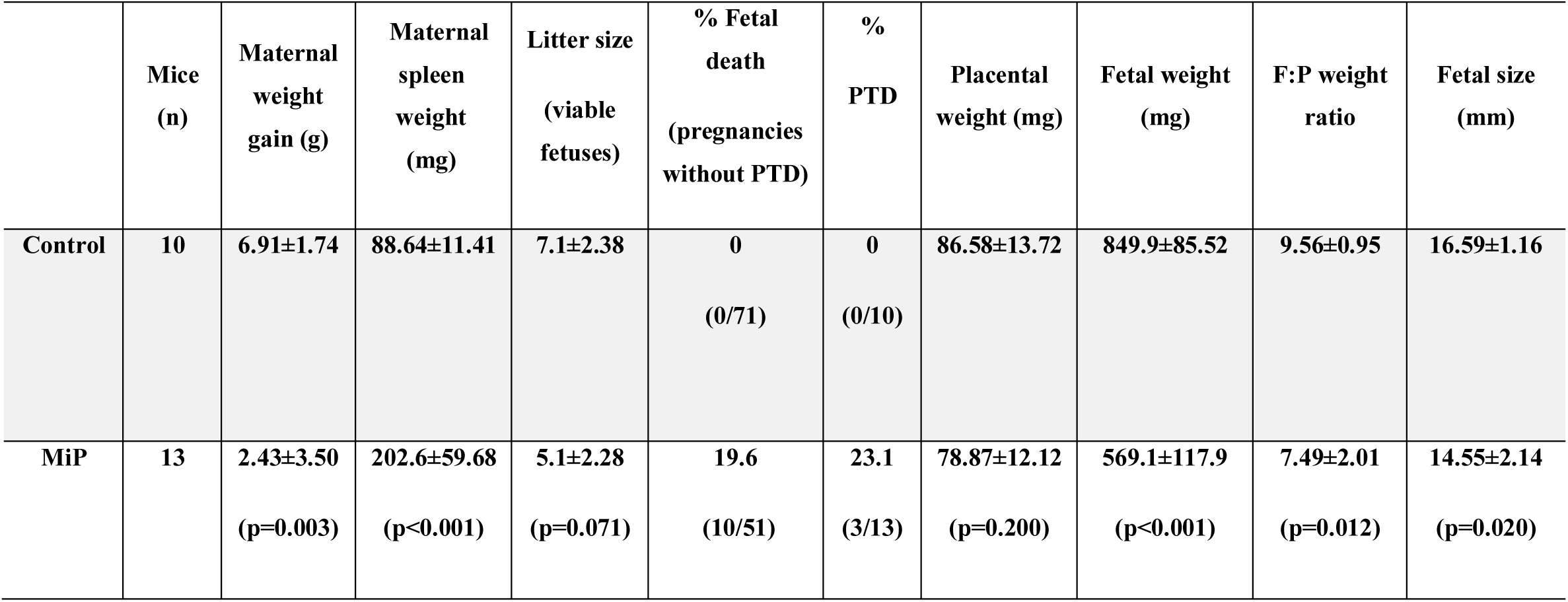
Summary of pregnancy parameters analyzed in *P. berghei* ANKA-infected mice and control mice at GD 18.5. Values are expressed as mean ± standard deviation.

|  | **Mice (n)** | **Maternal weight gain (g)** | **Maternal spleen weight (mg)** | **Litter size (viable fetuses)** | **% Fetal death (pregnancies without PTD)** | **% PTD** | **Placental weight (mg)** | **Fetal weight (mg)** | **F:P weight ratio** | **Fetal size (mm)** |
|---|---|---|---|---|---|---|---|---|---|---|
| **Control** | **10** | **6.91±1.74** | **88.64±11.41** | **7.1±2.38** | **0**<br>**(0/71)** | **0**<br>**(0/10)** | **86.58±13.72** | **849.9±85.52** | **9.56±0.95** | **16.59±1.16** |
| **MiP** | **13** | **2.43±3.50**<br>**(p=0.003)** | **202.6±59.68**<br>**(p<0.001)** | **5.1±2.28**<br>**(p=0.071)** | **19.6**<br>**(10/51)** | **23.1**<br>**(3/13)** | **78.87±12.12**<br>**(p=0.200)** | **569.1±117.9**<br>**(p<0.001)** | **7.49±2.01**<br>**(p=0.012)** | **14.55±2.14**<br>**(p=0.020)** |

In the acute sublethal endotoxemia model, as shown in Table 2, no changes in maternal weight gain (*P*=0.179), maternal spleen weight (*P*=0.307), litter size (*P*=0.866), placental weight (*P*=0.269), F:P weight ratio (*P*=0.085), and fetal size (*P*=0.078) were found. A reduction in fetal weight (*P*=0.039) was observed in LPS-injected dams, compared to the control group. Furthermore, acute sublethal endotoxemia promoted a PTD rate of 33.3% (5 of 15 LPS-injected dams had vaginally born dead fetuses between GD17.5, after LPS administration, and GD18.5, before euthanasia) and a fetal death rate of 4.1% (3 out of 74 fetuses of infected dams showing clear signs of non-viability).

**Table 2:** Summary of pregnancy parameters analyzed at GD 18.5, 24 hours after sublethal LPS challenge (LPS group) or PBS injection (control group). Values are expressed as mean ± standard deviation.

|  | **Mice (n)** | **Maternal weith gain (g)** | **Maternal spleen weight (mg)** | **Litter size (viable fetuses)** | **% Fetal death (pregnancies without PTD)** | **% PTD** | **Placental weight (mg)** | **Fetal weight (mg)** | **F:P weight ratio** | **Fetal size (mm)** |
|---|---|---|---|---|---|---|---|---|---|---|
| **Control** | **10** | **13.77±4.56** | **92.71±17.15** | **7.1±3.84** | **1.4<br>(1/71)** | **0<br>(0/10)** | **82.15±10.34** | **871.7±162.5** | **10.79±1.77** | **17.22±2.02** |
| **LPS** | **15** | **10.73±5.13<br>(p=0.179)** | **100.6±16.47<br>(p=0.307)** | **7.4±3.98<br>(p=0.866)** | **4.1<br>(3/74)** | **33.3<br>(5/15)** | **77.5±7.55<br>(p=0.269)** | **716.2±150.2<br>(p=0.039)** | **9.32±1.82<br>(p=0.085)** | **15.63±1.77<br>(p=0.078)** |

**Table 3.**
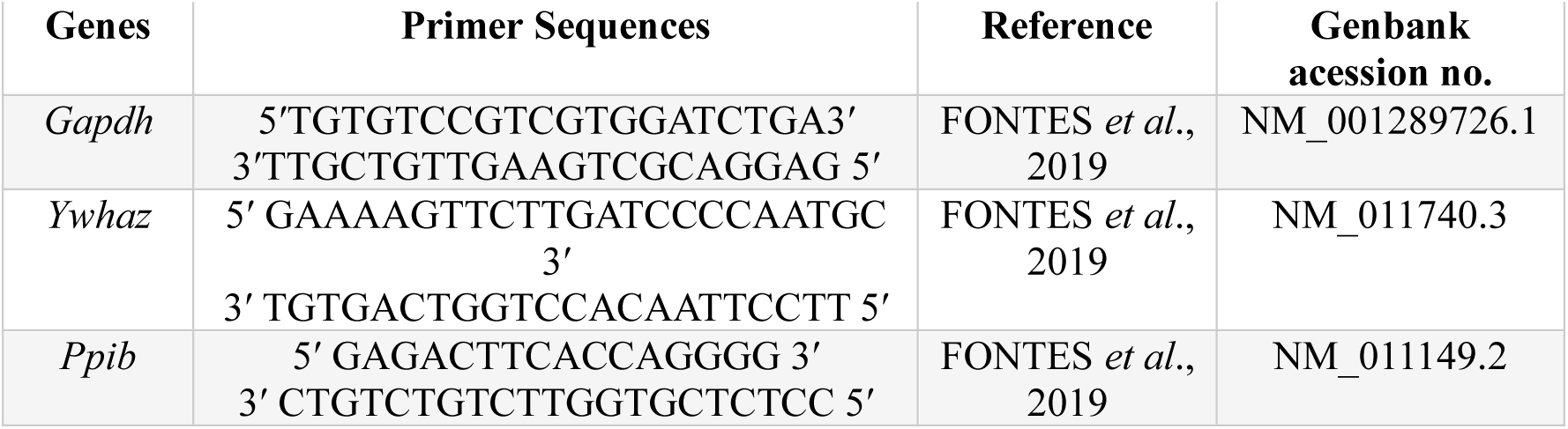
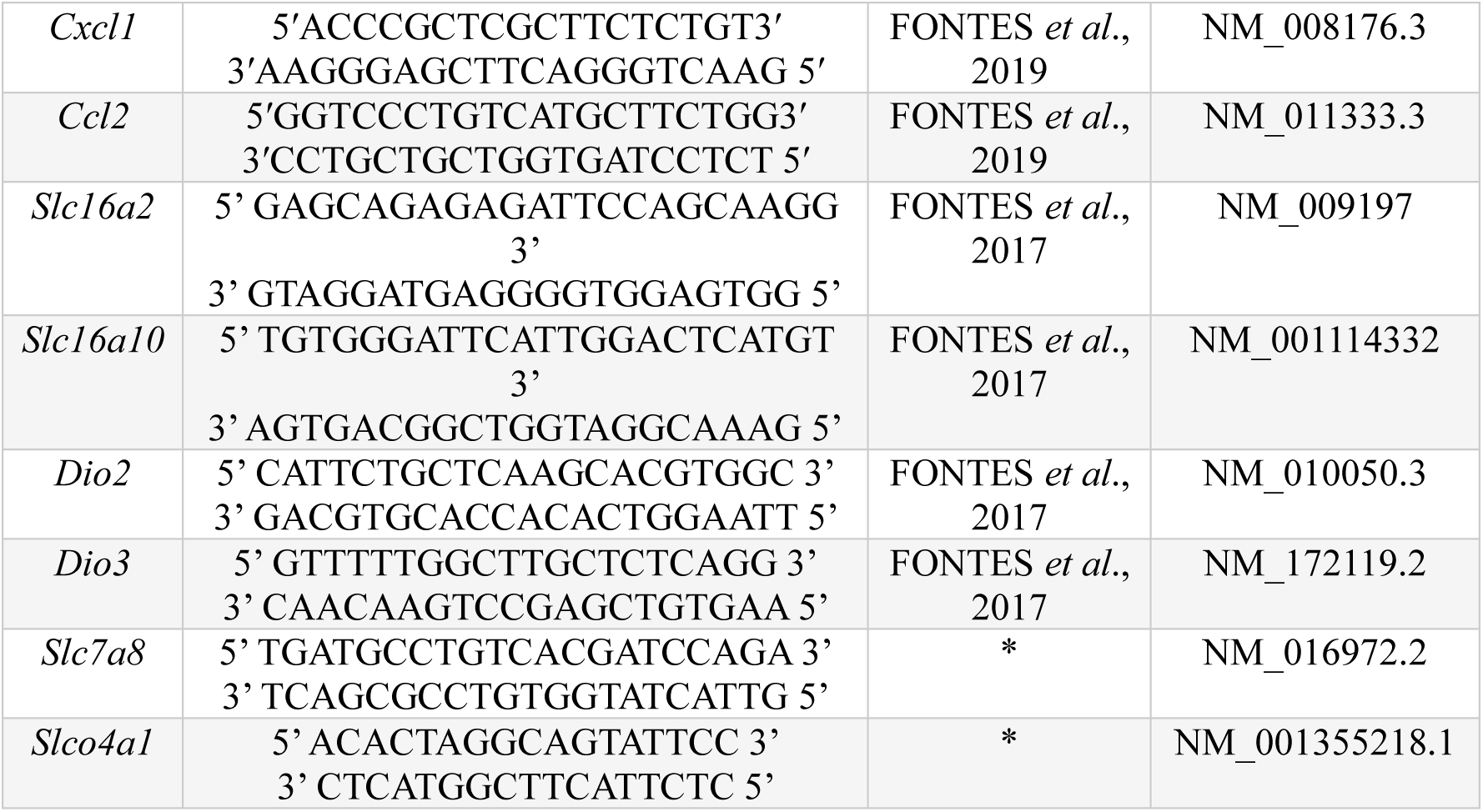
Primer’s sequences used in the study. *Primers designed with primer-BLAST (http://www.ncbi.nlm.gov/tools/primer-blast).

### MiP and acute sublethal endotoxemia impair hypothalamic-pituitary-thyroid (HPT) axis homeostasis

The effects of MiP and LPS exposure on the maternal HPT axis were evaluated by analyzing serum fT4 and TSH levels in the maternal serum (Fig. 2). The MiP infected dams had a reduction in serum fT4 (Fig. 2A, *P*=0.015) and an increase in TSH (Fig. 2B, *P*<0.001), compared to the control group. In the acute sublethal endotoxemia model, there were no changes in serum fT4 (Fig. 2C, *P*=0.315) but the TSH was increased in LPS-injected dams (Fig. 2D, *P*<0.001), compared to the control group.

**Figure 2.**
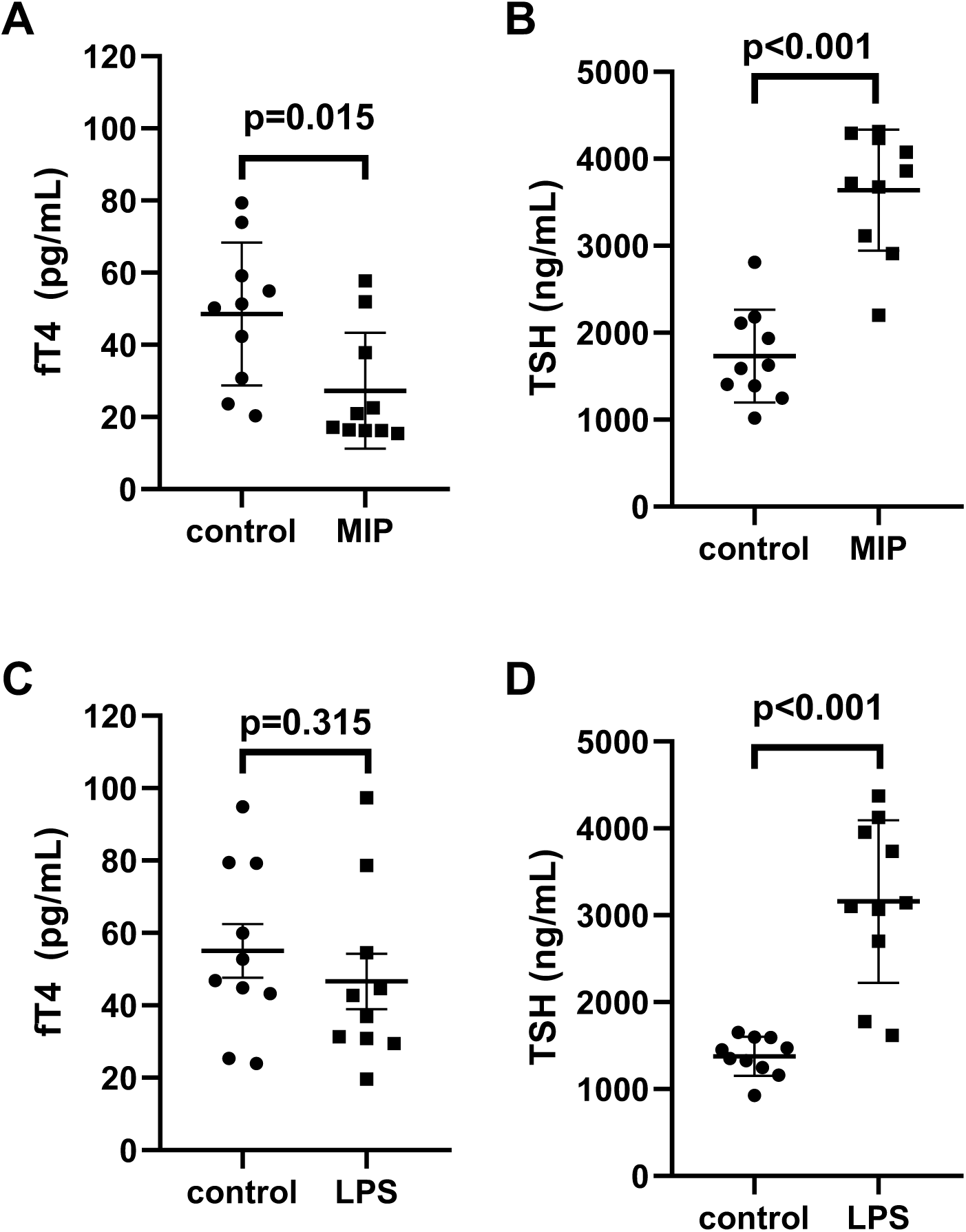
Maternal fT4 and TSH serum levels in MiP and LPS (24h) experimental models. **(A)** fT4 (MiP; n=10) **(B)** TSH (MiP; n=10); **(C)** fT4 (LPS 24h; n=10); **(D)** TSH (LPS 24h, n=10). Values are expressed as mean ± standard deviation. The Mann-Whitney test was employed for fT4 and TSH comparisons in both MiP and LPS models.

### MiP and LPS exposure alter the placental gene expression of inflammatory mediators, TH transporters and deiodinases

To investigate the effects of MiP and sublethal LPS exposure on placental TH metabolism, the mRNA expression of important placental TH transporters, *Slc16a2* (encoding MCT8); *Slc16a10* (MCT10); *Slc7a8* (LAT2); and *Slco4a1* (OATP4A1) as well as the iodothyronine deiodinases *Dio2* (D2) and *Dio3* (D3) were evaluated (Fig. 3 and 4). The mRNA expression of two important inflammatory markers, *Cxcl1* (CXCL1) and *Ccl2* (CCL2) associated with the onset of PTD, fetal death, and IUGR were also assessed. In the MiP model, there was a reduction in the placental mRNA expression of *Slc7a8* in both male and female offspring of infected dams (Fig. 3A-B, *P*=0.007 and *P*<0.001, respectively) and an upregulation in the placental mRNA expression of *Cxcl1* in both sexes, compared to control group (Fig. 3A-B, *P*=0.019 and *P*=0.022, respectively). We also observed a downregulation in placental mRNA expression of *Dio2* in male offspring of infected dams, compared to the control group (Fig. 3A, *P*<0.001). No significant differences were observed in the placental mRNA expression of *Dio2* in females, compared to control group (Fig. 3B, *P*=0.125) and also in the mRNA expression of *Slc16a2*, *Slc16a10*, *Slco4a1*, *Dio3* and *Ccl2*, in both male and female offspring of malaria-infected dams, compared to control group (Fig. 3A, *P*=0.126; 0.112; 0.288; 0.147 and 0.247, respectively and Fig. 3B, *P*=0.963; 0.340; 0.146; 0.131 and 0.868, respectively). In the LPS experimental model, there was an upregulation in the placental mRNA expression of *Ccl2* in male offspring of LPS-injected dams, compared to control group (Fig. 4A, *P*=0.049), whereas placental mRNA expression of *Cxcl1* was increased in female offspring of LPS-injected dams, compared to control group (Fig. 4B, *P*=0.011). No significant differences were observed in the placental mRNA expression of *Ccl2* in females (Fig. 4B, *P*=0.224) and *Cxcl1* in males (Fig. 4A, *P*=0.069), as well as in mRNA expression of *Slc16a2*, *Slc16a10*, *Slc7a8*, *Slco4a1*, *Dio2* and *Dio3*, in both male and female offspring of LPS-injected dams, compared to control group (Fig. 4A, *P*=0.061; 0.090; 0.118; 0.076; 0.989 and 0.588, respectively and Fig. 4B, *P*=0.361; 0.058; 0.250; 0.983; 0.635 and 0.263, respectively).

**Figure 3.**
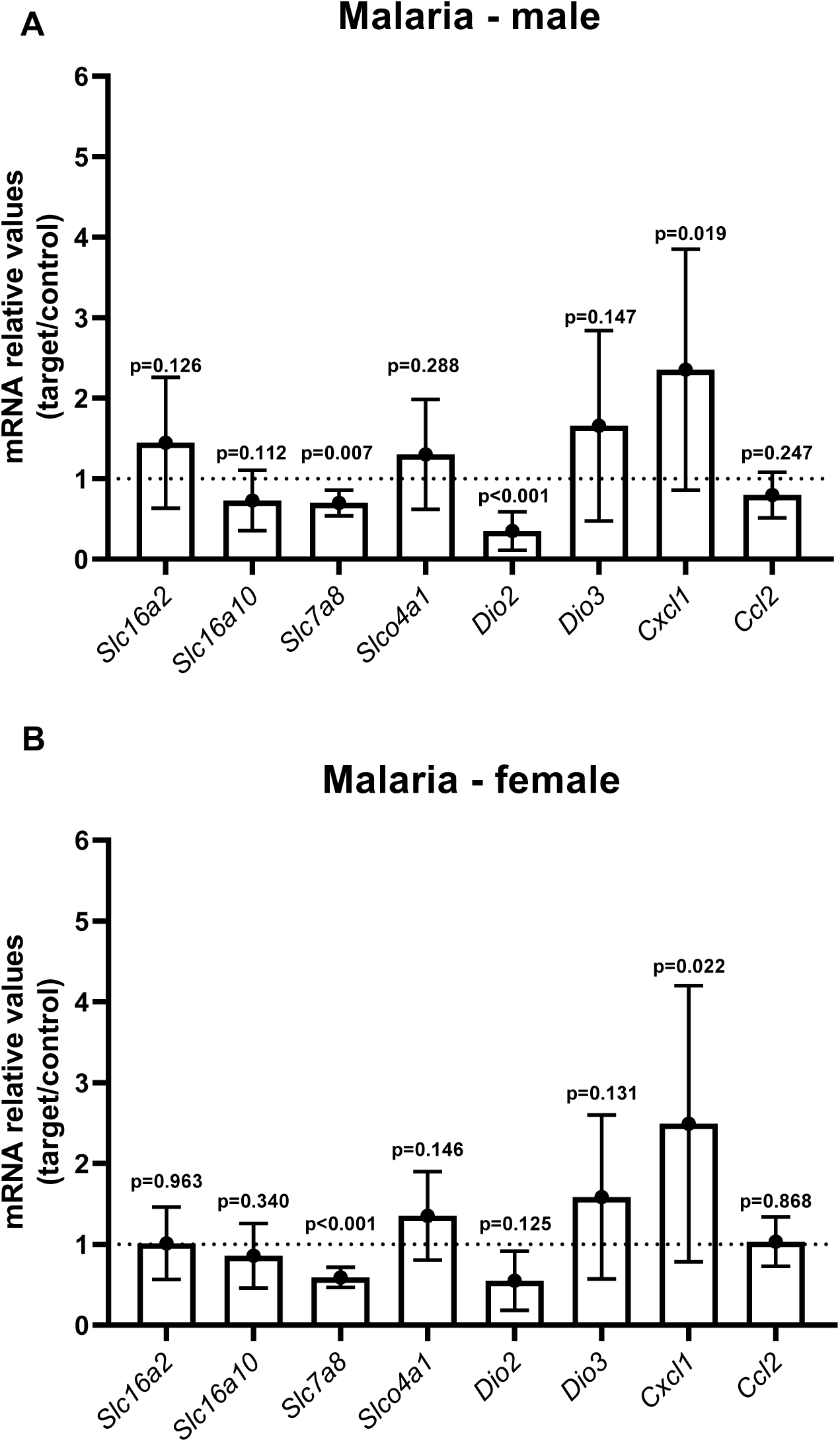
Malaria in pregnancy (MiP) regulates the expression of chemokines, TH transporters and deiodinases genes in the mouse placenta. mRNA levels of *Slc16a2, Slc16a10, Slc7a8, Slco4a1, Dio2, Dio3, Cxcl1* and *Ccl2* in the placenta of male **(A)** and female **(B)** offsprings. The broken line indicates the expression levels in the control group, which were set to one. Values are expressed as mean ± standard deviation. *n* = 10/group. Unpaired Students T-Tests were used in both analyzes of male and female placentas.

**Figure 4.**
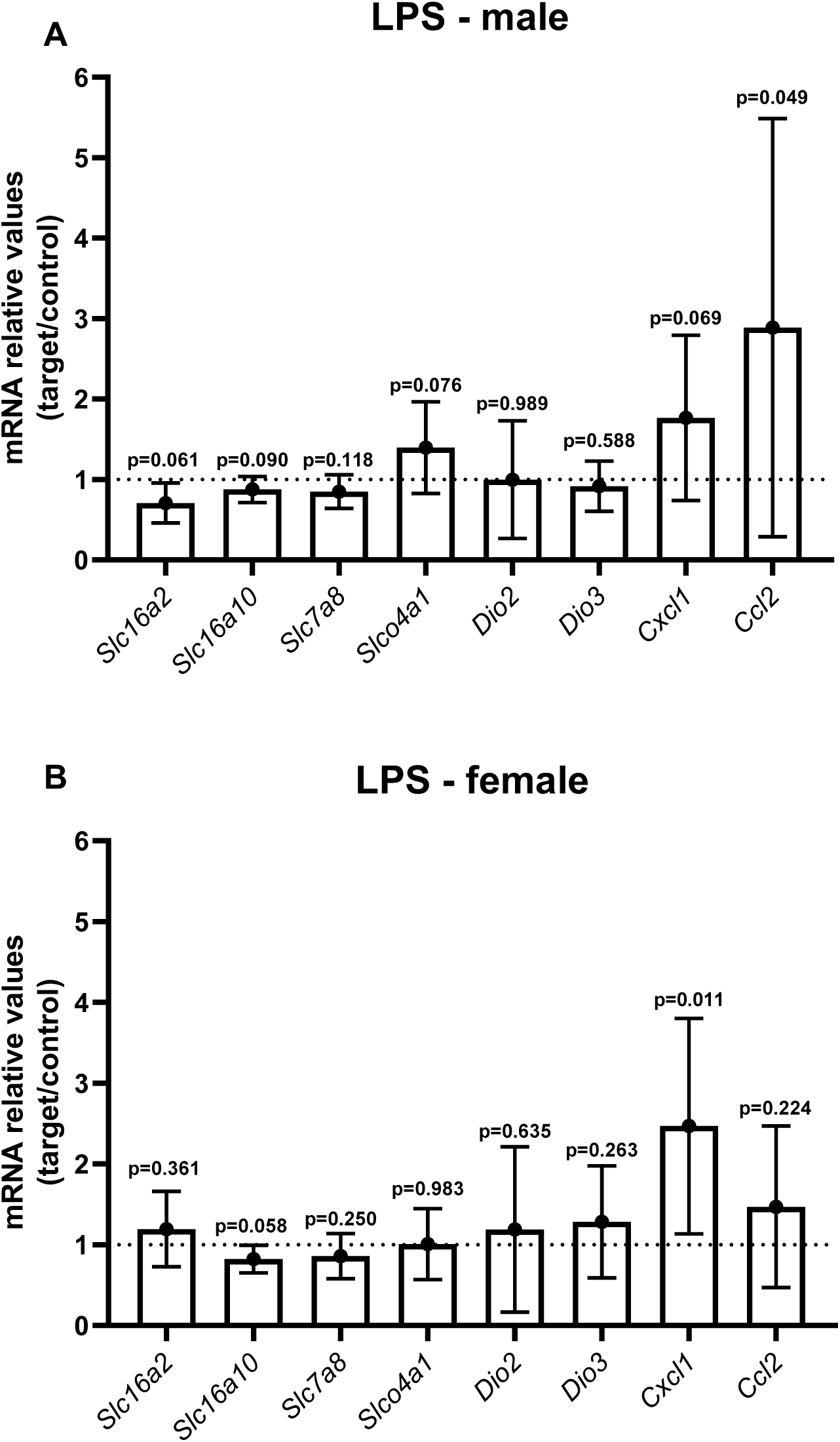
Sublethal LPS challenge (24h) regulates chemokine gene expression in the mouse placenta. mRNA levels of *Slc16a2, Slc16a10, Slc7a8, Slco4a1, Dio2, Dio3, Cxcl1* and *Ccl2* in the placenta of male **(A)** and female **(B)** offspring. The broken line indicates the expression levels in the control group, which were set to one. Values are expressed as mean ± standard deviation. *n* = 10/group. Unpaired Students T-Tests were used in both analyzes of male and female placentas.

### Malaria in pregnancy and LPS exposure modulate protein expression of TH transporters in the mouse placenta

The protein expression of MCT8, MCT10, LAT2 and OATP4A1 was assessed in both experimental models, given their role in the transport of iodothyronines in various tissues, including the human and murine placenta. In MiP, there was an upregulation of placental MCT8 expression in the Lz of both male and female offspring of infected dams, compared to the control group (Fig. 5A-D and 5E-H, *P*=0.028 and *P*=0.001, respectively). Furthermore, the expression of MCT8 was also increased in the placental Jz of male offspring of *Plasmodium*-infected mice, compared to the control group (Fig. 5 I-L, *P*=0.008). No significant differences were observed in the expression of MCT8 in the Jz of female offspring, compared to the control group (Fig. 5M-P, *P*=0.062). In the LPS experimental model, there was an upregulation of MCT8 only in the placentas of male offspring of LPS-injected dams, in both the Lz (Fig. 6A-D, *P*=0.001) and the Jz (Fig. 6I-L, *P*=0.001). No significant differences were observed in the expression of MCT8 in the Lz and Jz of female offspring, compared to the control group (Fig. 6E-H, *P*=0.637 and Fig. 6M-P, *P*=0.973, respectively).

**Figure 5.**
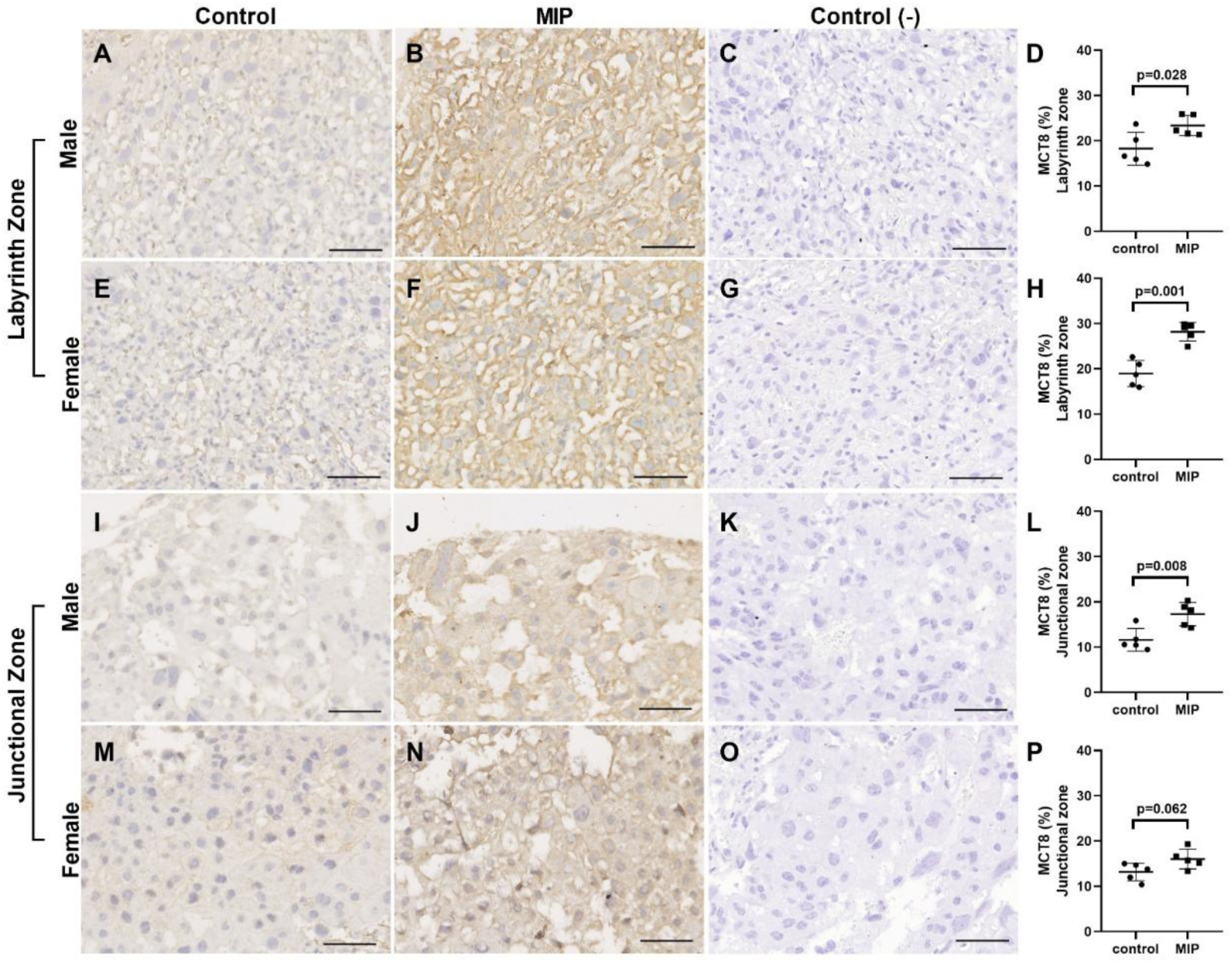
Malaria in pregnancy (MiP) regulates MCT8 protein expression in the mouse placenta. Representative photomicrographies of immunohistochemistry and semiquantitative analysis of MCT8 in the labyrinth zone (**A-D:** control and malaria-infected male offspring; **E-H:** control and malaria-infected female offspring) and junctional zone (**I-L:** control and malaria-infected male offspring; **M-P:** control and malaria-infected female offspring). The graph represents the % of stained cells per area of viable tissue. The brown color indicates immunolocalization of the antibody on cell membranes containing the MCT8 antigen. Control (-) represents the negative control (without primary antibody). Values are expressed as mean ± standard deviation. *n* = 5/group. Unpaired Students T-Tests were used in both analyzes of male and female placentas. Scale bar = 50 µm.

**Figure 6.**
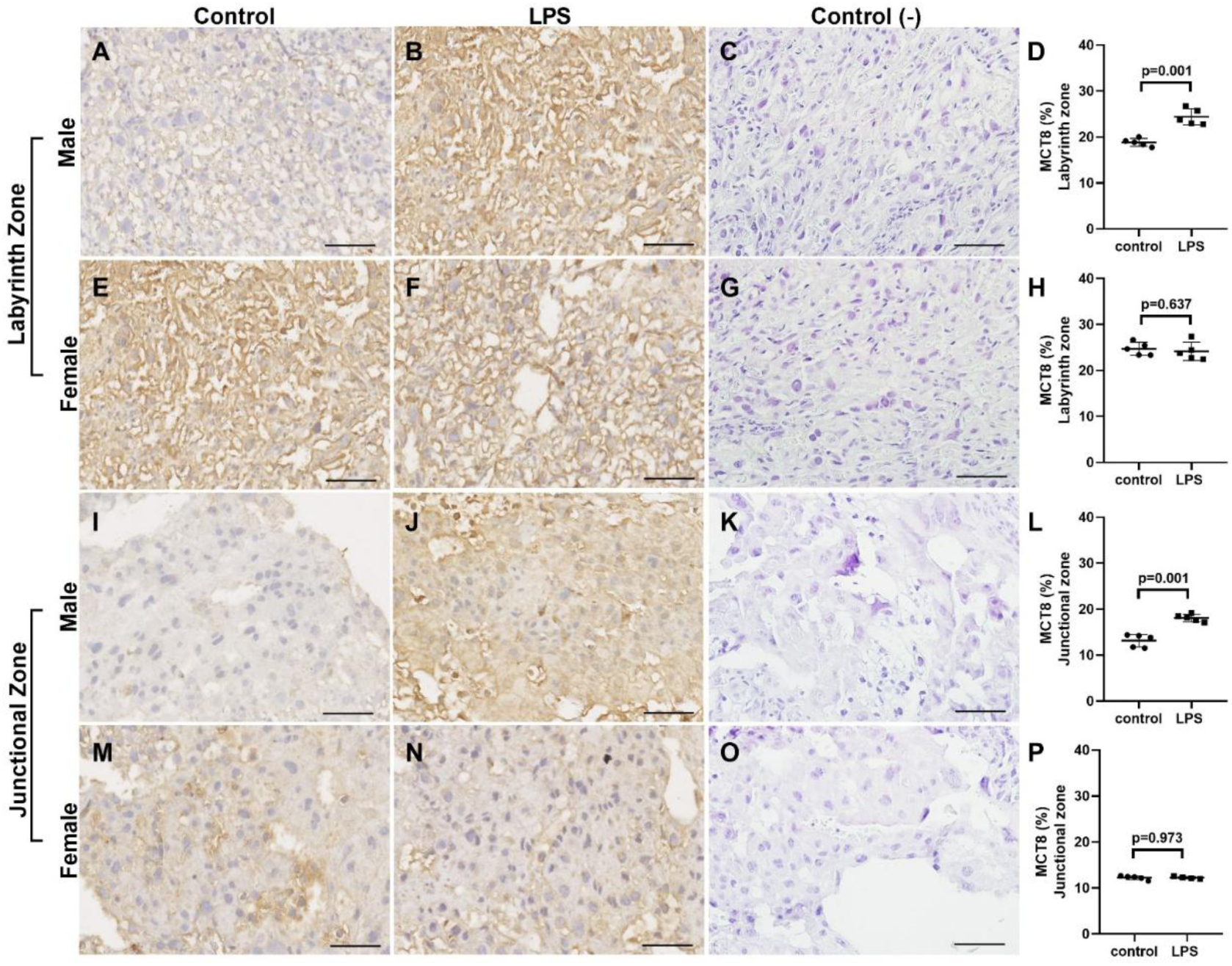
Sublethal LPS challenge (24h) regulates MCT8 protein expression in the mouse placenta. Representative photomicrographies of immunohistochemistry and semiquantitative analysis of MCT8 in the labyrinth zone (**A-D:** control and LPS-injected male offspring; **E-H:** control and LPS-injected female offspring) and junctional zone (**I-L:** control and LPS-injected male offspring; **M-P:** control and LPS-injected female offspring). The graph represents the % of stained cells per area of viable tissue. The brown color indicates immunolocalization of the antibody on cell membranes containing the MCT8 antigen. Control (-) represents the negative control (without primary antibody). Values are expressed as mean ± standard deviation. Values are expressed as mean ± standard error of the mean. *n* = 5/group. Unpaired Students T-Tests were used in both analyzes of male and female placentas. Scale bar = 50 µm.

MCT10 expression was downregulated in the placental labyrinth zone of both male and female offspring of malaria-infected dams, compared to the control group (Fig. 7A-D and 7E-H, *P*=0.001 and *P*<0.001, respectively). In turn, MCT10 expression was upregulated in the placental junctional zone of both male and female offspring of malaria-infected dams (Fig. 7I-L and 7M-P, *P*<0.001). Similar to what was observed in MiP experimental model, MCT10 expression was downregulated in the placental labyrinth zone of both male and female offspring of LPS-injected dams (Fig. 8A-D and 8E-H, *P*<0.001). On the other hand, MCT10 expression was upregulated in the placental junctional zone of both male and female offspring of LPS-injected dams (Fig. 8I-L and 8M-P, *P*<0.001).

**Figure 7.**
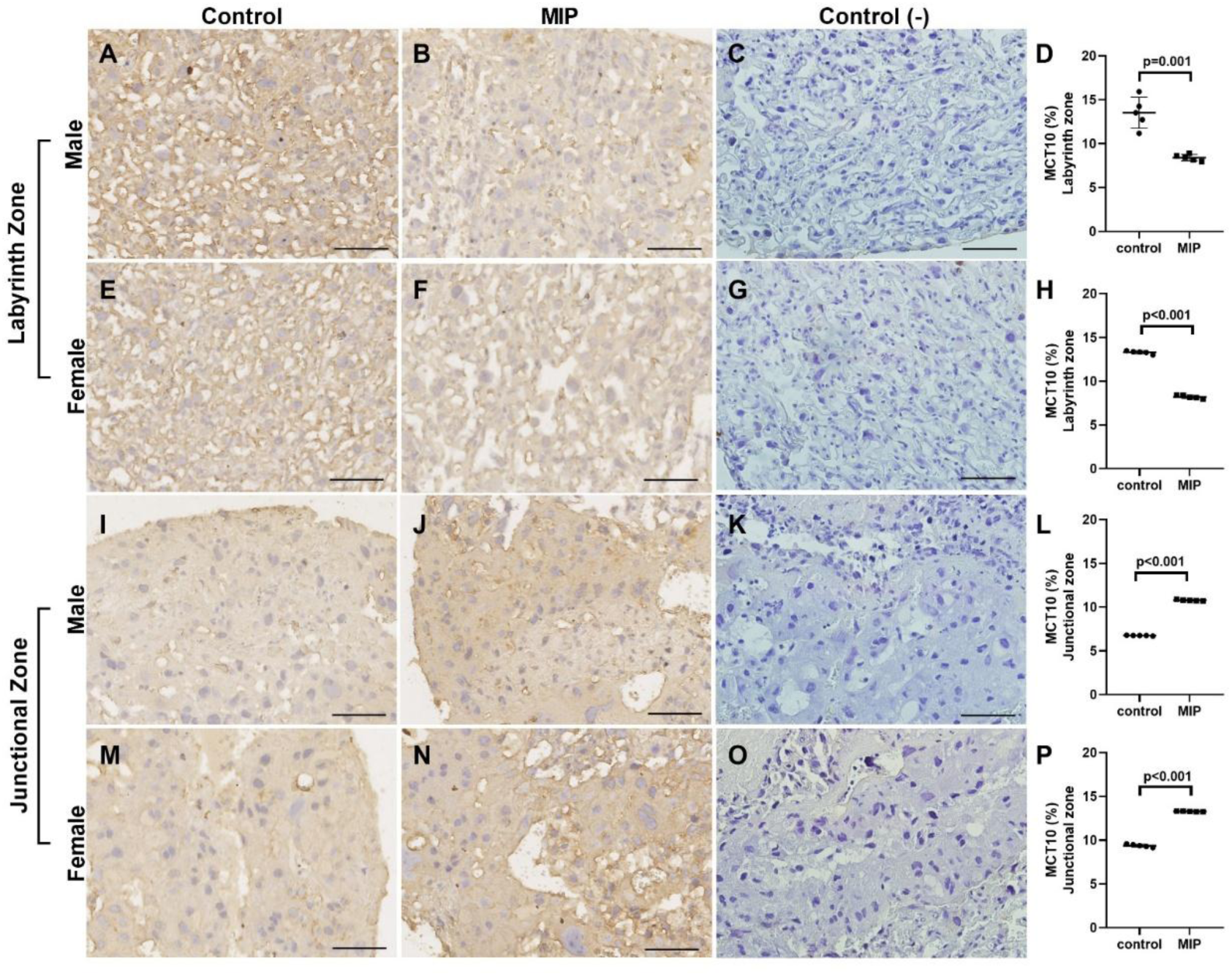
Malaria in pregnancy (MiP) regulates MCT10 protein expression in the mouse placenta. Representative photomicrographies of immunohistochemistry and semiquantitative analysis of MCT10 in the labyrinth zone (**A-D:** control and malaria-infected male offspring; **E-H:** control and malaria-infected female offspring) and junctional zone (**I-L:** control and malaria-infected male offspring; **M-P:** control and malaria-infected female offspring). The graph represents the % of stained cells per area of viable tissue. The brown color indicates immunolocalization of the antibody on cell membranes containing the MCT10 antigen. Control (-) represents the negative control (without primary antibody). Values are expressed as mean ± standard deviation. *n* = 5/group. Unpaired Students T-Tests were used in both analyzes of male and female placentae. Scale bar = 50 µm.

**Figure 8.**
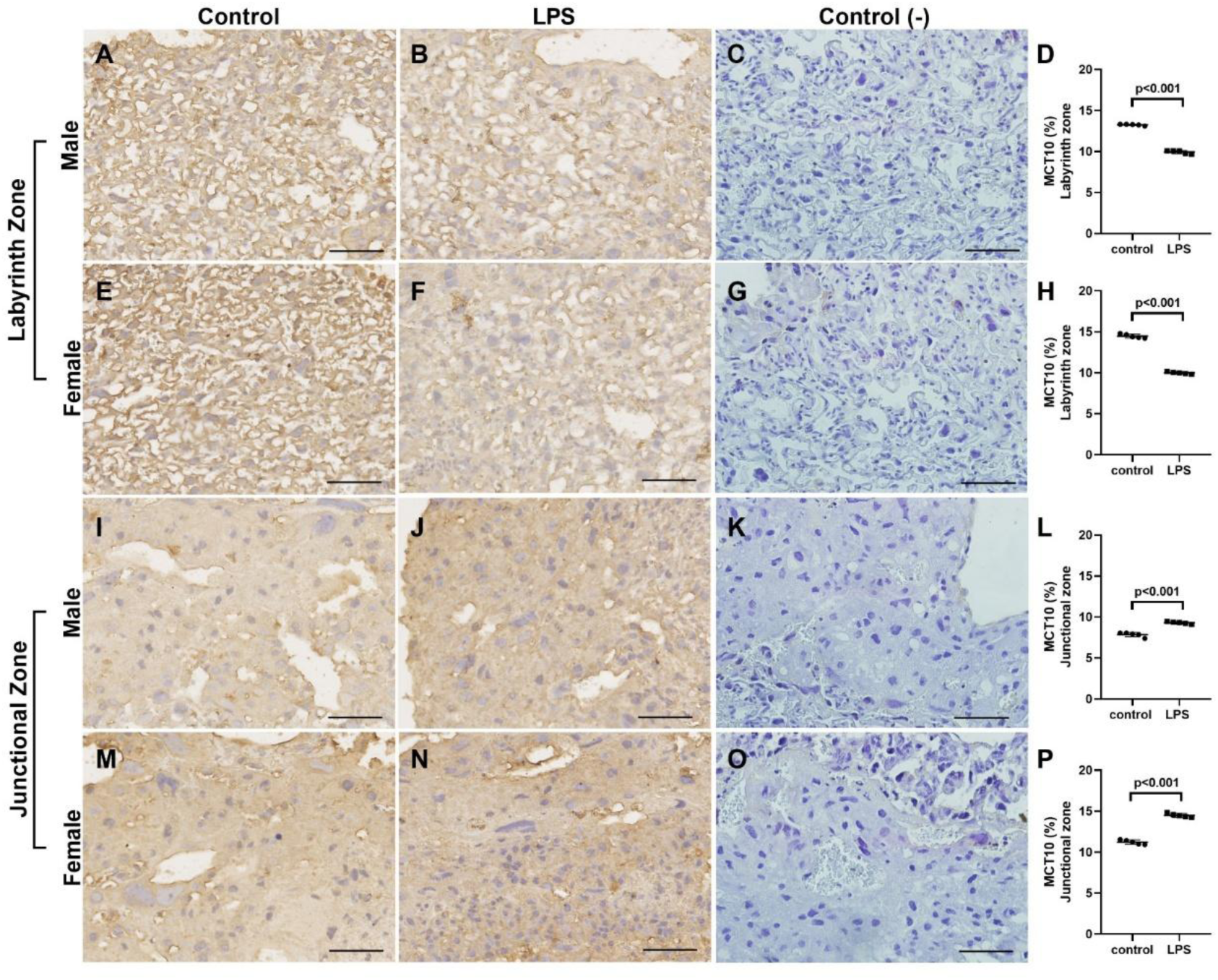
Sublethal LPS challenge (24h) regulates MCT10 protein expression in the mouse placenta. Representative photomicrographies of immunohistochemistry and semiquantitative analysis of MCT10 in the labyrinth zone (**A-D:** control and LPS-injected male offspring; **E-H:** control and LPS-injected female offspring) and junctional zone (**I-L:** control and LPS-injected male offspring; **M-P:** control and LPS-injected female offspring). The graph represents the % of stained cells per area of viable tissue. The brown color indicates immunolocalization of the antibody on cell membranes containing the MCT10 antigen. Control (-) represents the negative control (without primary antibody). Values are expressed as mean ± standard deviation. *n* = 5/group. Unpaired Students T-Tests were used in both analyzes of male and female placentae. Scale bar = 50 µm.

LAT2 expression was upregulated only in male offspring of malaria-infected dams, in both placental labyrinth and junctional zones (Fig. 9A-D and 9I-L, *P*<0.001 and *P*=0.001, respectively), while it was also upregulated in the placental Lz of male offspring of LPS-injected dams (Fig. 10A-D, *P*<0.001). No significant differences were observed in the expression of LAT2 in the Lz and Jz of female offspring of malaria-infected dams, compared to control group (Fig. 9E-H, *P*=0.493 and Fig. 9M-P, *P*=0.278, respectively), while no significant differences were observed either in the Lz of females (Fig. 10E-H, *P*=0.640) and in the Jz of male and female offspring of LPS-injected dams (Fig. 10I-L, *P*=0.797 and Fig. 10M-P, *P*=0.089, respectively).

**Figure 9.**
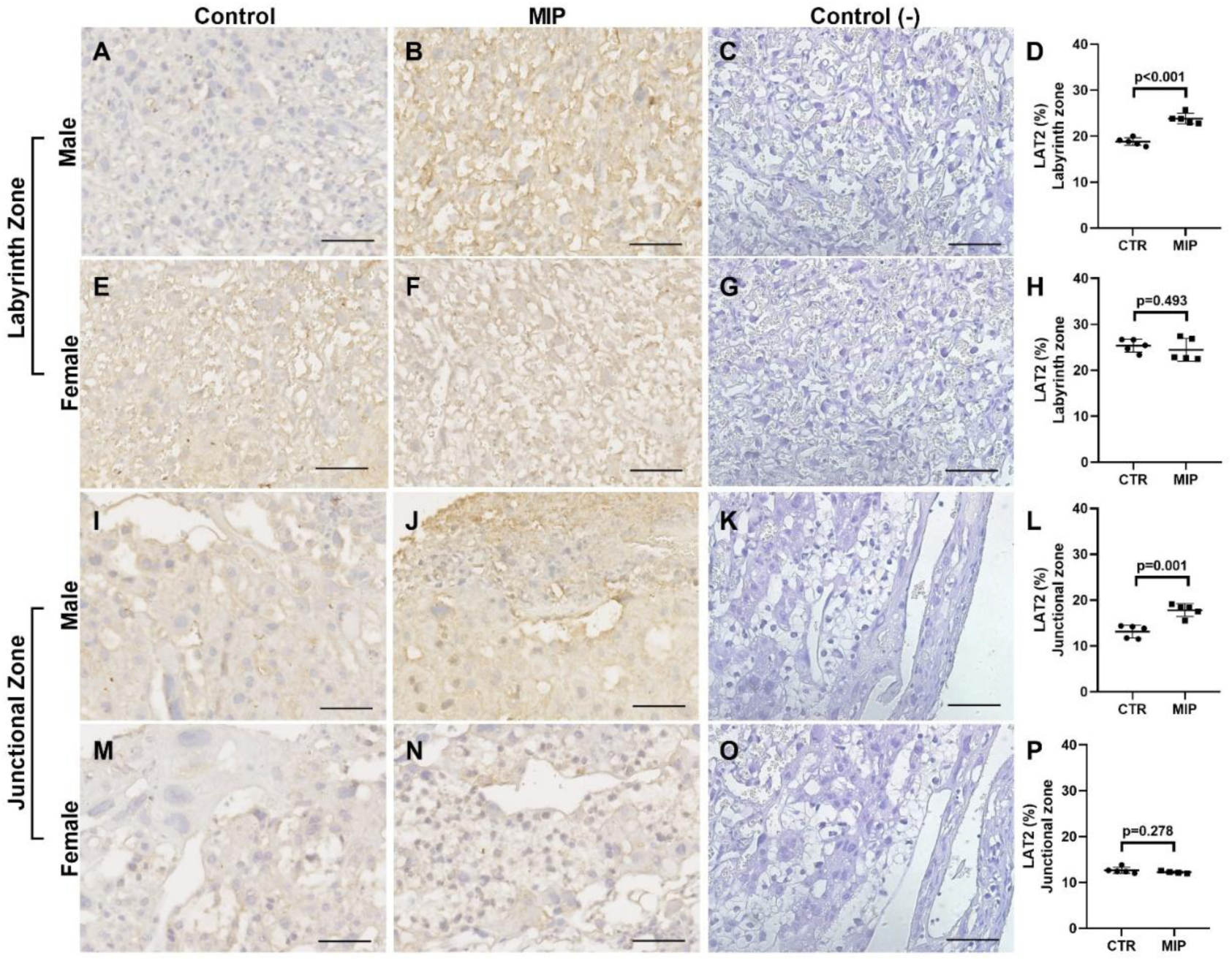
Malaria in pregnancy (MiP) regulates LAT2 protein expression in the mouse placenta. Representative photomicrographies of immunohistochemistry and semiquantitative analysis of LAT2 in the labyrinth zone (**A-D:** control and malaria-infected male offspring; **E-H:** control and malaria-infected female offspring) and junctional zone (**I-L:** control and malaria-infected male offspring; **M-P:** control and malaria-infected female offspring). The graph represents the % of stained cells per area of viable tissue. The brown color indicates immunolocalization of the antibody on cell membranes containing the LAT2 antigen. Control (-) represents the negative control (without primary antibody). Values are expressed as mean ± standard deviation. *n* = 5/group. Unpaired Students T-Tests were used in both analyzes of male and female placentae. Scale bar = 50 µm.

**Figure 10.**
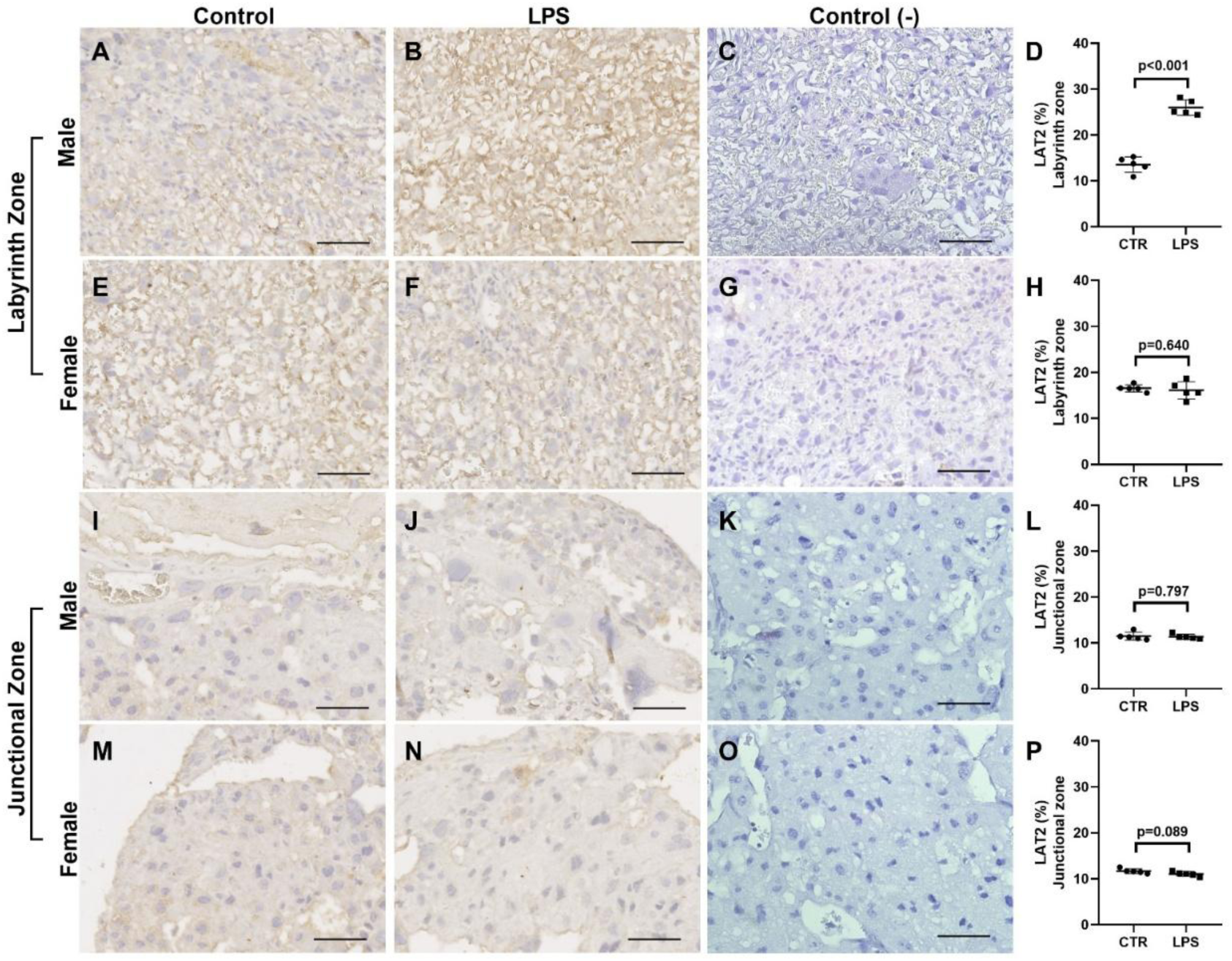
Sublethal LPS challenge (24h) regulates LAT2 protein expression in the mouse placenta. Representative photomicrographies of immunohistochemistry and semiquantitative analysis of LAT2 in the labyrinth zone (**A-D:** control and LPS-injected male offspring; **E-H:** control and LPS-injected female offspring) and junctional zone (**I-L:** control and LPS-injected male offspring; **M-P:** control and LPS-injected female offspring). The graph represents the % of stained cells per area of viable tissue. The brown color indicates immunolocalization of the antibody on cell membranes containing the LAT2 antigen. Control (-) represents the negative control (without primary antibody). Values are expressed as mean ± standard deviation. *n* = 5/group. Unpaired Students T-Tests were used in both analyzes of male and female placentae. Scale bar = 50 µm.

In MiP experimental model, similar to what was observed in LAT2 expression, OATP4A1 expression was upregulated only in male offspring of malaria-infected dams, in both placental labyrinth and junctional zones (Fig. 11A-D and 11I-L, *P*<0.001 and *P*=0.001, respectively), while it was upregulated in the placental Lz of both male and female offspring of LPS injected-dams, compared to control group (Fig. 12A-D and 12E-H, *P*=0.009 and *P*<0.001, respectively). No significant differences were observed in the expression of OATP4A1 in the Lz and Jz of female offspring of malaria-infected dams (Fig. 11E-H, *P*=0.493 and Fig. 11M-P, *P*=0.670, respectively), while no significant differences were observed either in the Jz of male and female offspring of LPS-injected dams, (Fig. 12I-L, *P*=0.399 and Fig. 12M-P, *P*=0.390, respectively).

**Figure 11.**
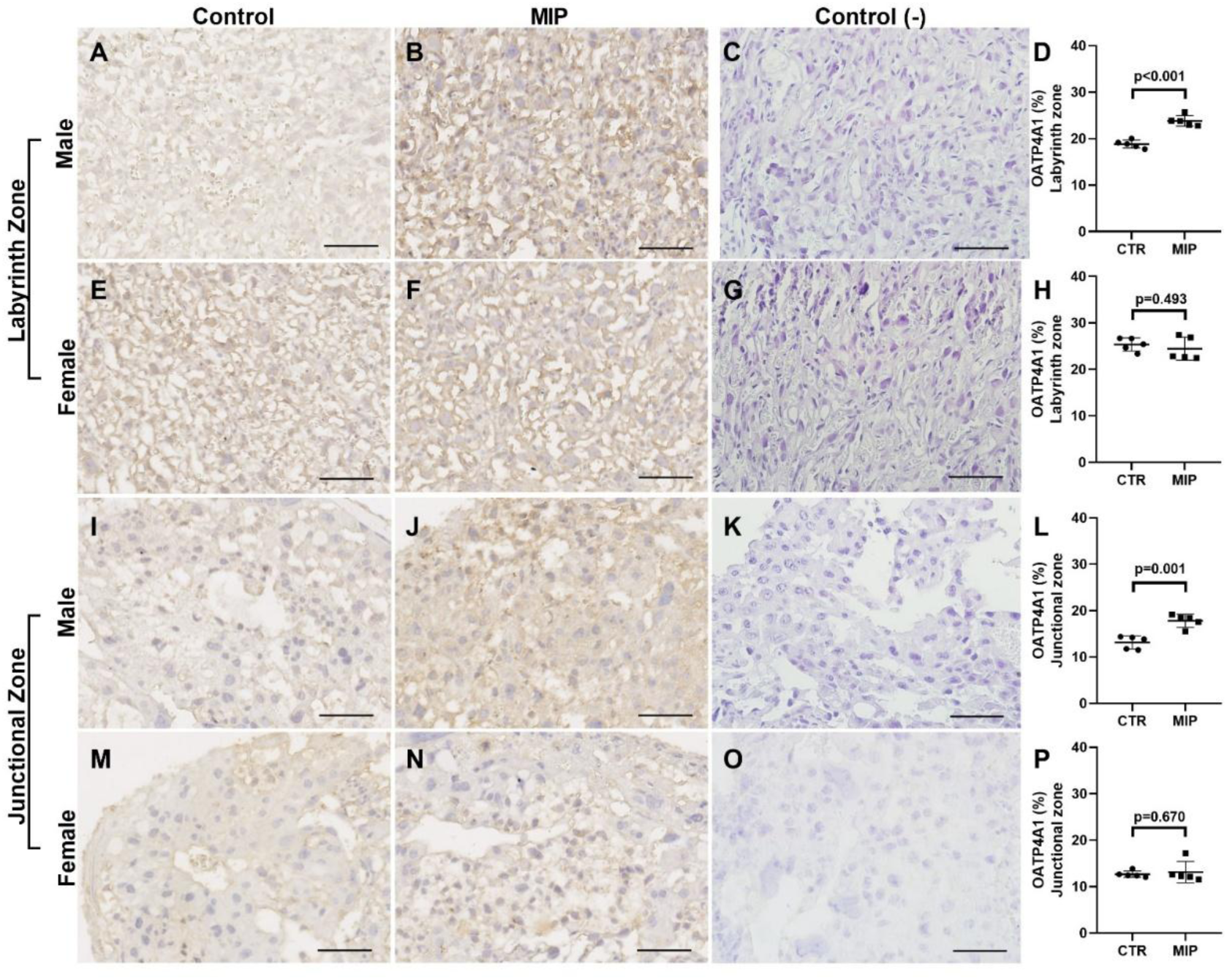
Malaria in pregnancy (MiP) regulates OATP4A1 protein expression in the mouse placenta. Representative photomicrographies of immunohistochemistry and semiquantitative analysis of OATP4A1 in the labyrinth zone (**A-D:** control and malaria-infected male offspring; **E-H:** control and malaria-infected female offspring) and junctional zone (**I-L:** control and malaria-infected male offspring; **M-P:** control and malaria-infected female offspring). The graph represents the % of stained cells per area of viable tissue. The brown color indicates immunolocalization of the antibody on cell membranes containing the OATP4A1 antigen. Control (-) represents the negative control (without primary antibody). Values are expressed as mean ± standard deviation. *n* = 5/group. Unpaired Students T-Tests were used in both analyzes of male and female placentae. Scale bar = 50 µm.

**Figure 12.**
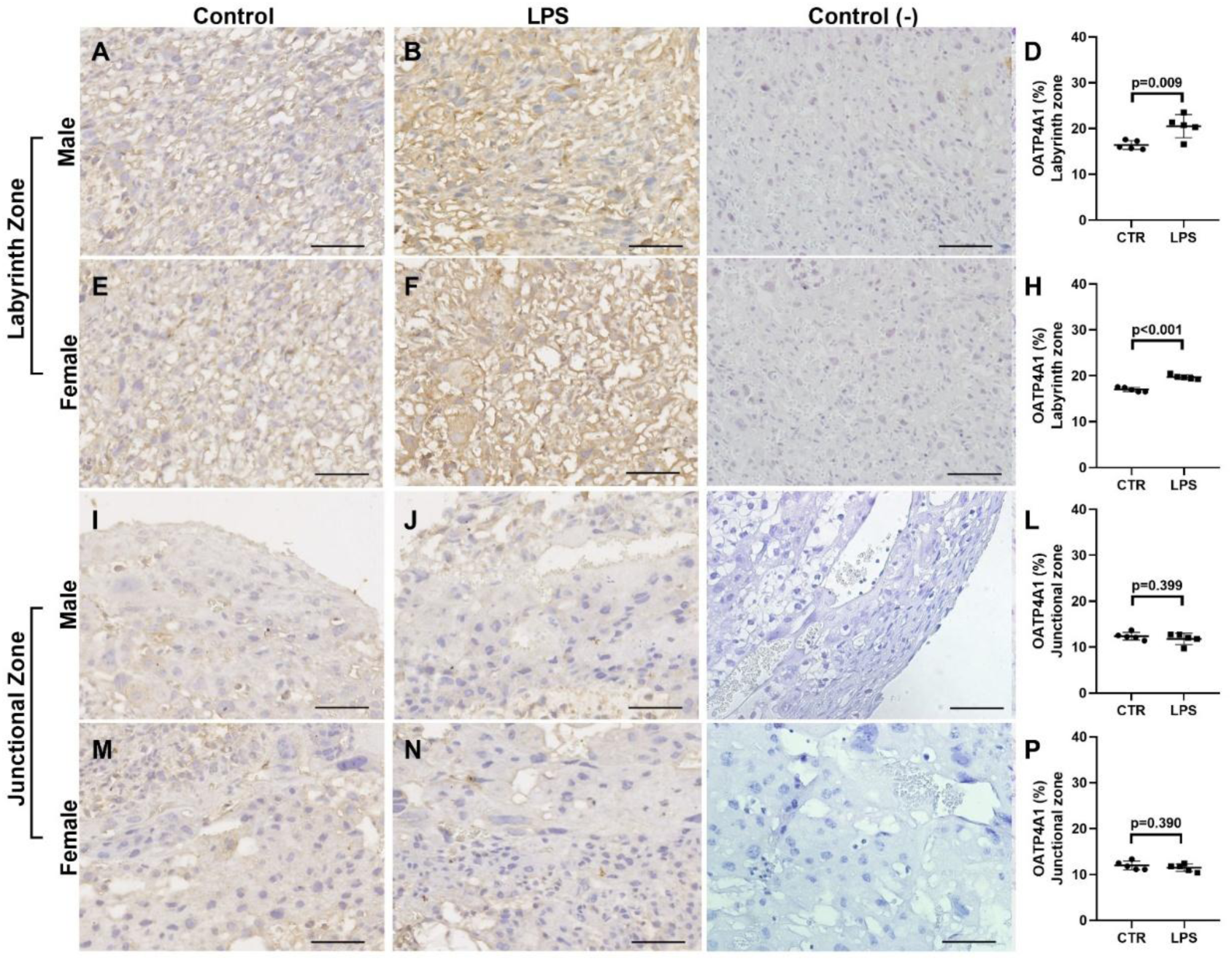
Sublethal LPS challenge (24h) regulates OATP4A1 protein expression in the mouse placenta. Representative photomicrographies of immunohistochemistry and semiquantitative analysis of OATP4A1 in the labyrinth zone (**A-D:** control and LPS-injected male offspring; **E-H:** control and LPS-injected female offspring) and junctional zone (**I-L:** control and LPS-injected male offspring; **M-P:** control and LPS-injected female offspring). The graph represents the % of stained cells per area of viable tissue. The brown color indicates immunolocalization of the antibody on cell membranes containing the OATP4A1 antigen. Control (-) represents the negative control (without primary antibody). Values are expressed as mean ± standard deviation. *n* = 5/group. Unpaired Students T-Tests were used in both analyzes of male and female placentae. Scale bar = 50 µm.

## DISCUSSION

This study reports that MiP and sublethal endotoxemia induced by acute LPS challenge impair maternal thyroid function and the expression of important placental TH transporters and deiodinases, in addition to promoting an inflammatory process at the maternal-fetal interface, which may be associated with fetak growth restriction observed in the MiP model and with isolated reduction in fetal weight observed after LPS challenge (Poovassery & Moore, 2006; Neres *et al*., 2008; Poovassery *et al*., 2009; Fontes *et al*., 2019; Reginatto *et al*., 2021). It is important to note that our model of MiP has been shown to recapitulate features of placental malaria in mice, including placental sequestration of infected erythrocytes, one of the hallmarks of MiP pathophysiology (Neres *et al*., 2008; De Oca *et al*., 2013; Fontes *et al*., 2019). Key features of placental malaria include fetal death, decreased fetal weight/ fetal size, decreased placental efficiency, reduced maternal weight gain and increased maternal spleen weight, all of which were observed in our MiP model. We have previously demonstrated that infection of C57BL/6 mice with *P. berghei* ANKA recapitulates placental sequestration of infected erythrocytes, leading to the establishment of a pro-inflammatory microenvironment at the maternal-fetal interface (Fontes *et al*., 2019). This is also observed in other MiP models and is related to decreased placental efficiency and altered expression of transport systems (Neres *et al*., 2008; Cressman *et al*., 2014; Najjar *et al*., 2019; Fontes *et al*., 2019).

Likewise, bacterial infections in pregnancy are linked to adverse maternal-fetal outcomes, including IUGR, fetal death, PTD and neonatal neurological damage, primarily due to increased expression of pro-inflammatory markers, reduced placental efficiency from altered barrier morphology and altered expression and function of nutrient and protective transporter systems (Fowden *et al*., 2009; Hayward *et al*., 2016; Liu *et al*., 2016; Reginatto *et al*., 2021; Nardi *et al*., 2025). In our murine model, sublethal LPS exposure resulted in 33.3% PTD and lower fetal weight, but with limited fetal death rates, corroborating previous studies reporting higher PTD in the late gestation induced by bacterial infections (Toyama *et al*., 2015; Reginatto *et al*., 2021).

The average gestational length for C57BL/6 mice is 19.25 days (Murray *et al*., 2010). The GD18.5 time point was chosen for euthanasia because this time point is considered as term stage of pregnancy and because fetal growth and placental volume/circulation reached the peak of their development (Coan *et al*., 2004). Vaginally born dead fetuses between GD17.5 and GD18.5 were considered premature since this time point is consistent with well-established markers of PTD in mice (McCarthy *et al*., 2018). Therefore, all analyses were performed with dams that did not undergo preterm labor and with viable placentas and fetuses, without signs of fetal death.

Systemic infections and acute inflammation impair the HPT axis, leading to non-thyroidal illness syndrome (NTIS), a condition marked by a reduction in serum TSH and TH levels (Fontes *et al*., 2017). The effects of severe malaria on HPT axis homeostasis and thyroid function in humans indicate that gestational malaria can promote NTIS, due to inappropriately low serum TSH levels (Davis *et al*., 1990). However, in our MiP experimental model, serum analyses indicate the establishment of primary hypothyroidism, since we observed a reduction in serum fT4 levels in *P. berghei* ANKA-infected dams, as well as an increase in serum TSH levels in both malaria-infected and LPS-injected dams. Primary hypothyroidism during pregnancy may originate primarily from placental hormonal production, especially estrogens, directly affecting thyroid function and reducing TH bioavailability (low fT4 and increased serum TSH and thyroglobulin levels) (Mallawa Kankanamalage *et al*., 2021).

Furthermore, hypothyroidism affects the integrity of the placental barrier, promoting morphological and vascular changes in the placental Lz, which are associated with IUGR (Serakides *et al*., 2012). Although literature evidence demonstrates that LPS administration in non-pregnant mice leads to the development of NTIS (Boelen *et al*., 2004), it is important to highlight that, in our experimental model, the different LPS dose and route of administration, as well as placental production of estrogens, pro-angiogenic and growth factors (Mallawa Kankanamalage *et al*., 2021), may explain the onset of primary subclinical hypothyroidism, a hypothesis that requires further investigation.

Hypothyroidism promotes the establishment of a pro-inflammatory placental environment and impairs extravillous trophoblasts migration and placentation (Silva *et al*., 2014). TH inhibit the activation of the NLRP3 inflammasome (De Luca *et al*., 2021) and, in rats, low serum TH levels stimulate the placental activation of this inflammasome (Santos *et al*., 2023), which are related to preeclampsia and IUGR(Menkhorst *et al*., 2023). This evidence demonstrates the importance of TH for trophoblast differentiation and proliferation, as well as vascular remodelling and integrity of the placental barrier, which, together with changes in placental transport and metabolism of TH, may be related to IUGR and reduced placental efficiency (Appiah Adu-Gyamfi *et al*., 2020).

In the MiP experimental model, we observed an increase in the placental *Cxcl1* mRNA expression in both male and female offspring of malaria-infected dams. In turn, *Cxcl1* mRNA expression was upregulated only in the placenta of female offspring of LPS-injected dams. In contrast, placental *Ccl2* mRNA expression was upregulated only in the male offspring of LPS-injected dams, with no changes in MiP model. CXCL1 promotes a pro-inflammatory environment associated with PTD and IUGR, and CCL2 induces leukocyte recruitment and inflammation in late gestation, also associated with IUGR in mice (Peltier, 2003; Loubière *et al*., 2010; Inoue *et al*., 2016). In addition, CCL2 activation by ligands of the innate immunity receptor NOD-1, in the murine placenta, promotes IUGR (Inoue *et al*., 2016). It is possible that, in both models, a combination of maternal thyroid dysfunction, overt after MiP and subclinical after LPS challente, and stimulation of the placental immune system contributes to the observed outcomes.

Since maternal-fetal transfer of TH depends on its placental transport and metabolism (Visser, 2016; Appiah Adu-Gyamfi *et al*., 2020), we set out to investigate the mRNA expression of the main TH transporters and iodothyronine deiodinases in the placenta. The TH transporters herein analyzed are mainly expressed in the syncytiotrophoblast, especially MCT8 and MCT10 (Loubière *et al*., 2010). We observed a reduction in *Slc7a8* (LAT2) mRNA expression in the placentas of male and female offspring from malaria-infected mice. LAT2 is a heterodimeric protein that transports neutral amino acids and iodothyronines, including T3 (Zevenbergen *et al*., 2015). However, C57BL/6 *Slc7a8-null* mice exhibit normal growth serum iodothyronine levels, suggesting functional redundancy among transporters (Braun *et al*., 2011).

In contrast to what was observed regarding mRNA expression, IHC analyses demonstrated an increase in the placental LAT2 expression, in both the labyrinth and the junctional zones, only in the male offspring of *Plasmodium*-infected dams. This same upregulation in LAT2 expression was observed only in the placental labyrinth of male offspring from dams injected with LPS. It has been previously demonstrated that BALB/c mice infected with *P. berghei* ANKA exhibit reduced expression of other amino acid transporters associated with IUGR (Najjar *et al*., 2019). Since system L amino acid transporters, specially LAT2, are highly expressed in the syncytiotrophoblast of human third trimester placentas (Gaccioli *et al*., 2015), and also in the labyrinth and junctional zones of rodent placentas (Owaydhah *et al*., 2021), it is possible that the differences observed between LAT2 mRNA and protein expression reflect a compensatory mechanism associated with a potential reduction in placental transport of TH and amino acids, a hypothesis that also requires further investigation.

MCT8 is a specific transporter of iodothyronines (Friesema *et al*., 2003). We evaluated the mRNA and protein expression of this transporter in the placentas of both experimental models. Although *Slc16a2* mRNA was unchanged in both models, placental MCT8 protein expression was upregulated in the Lz and Jz of male placentas in both models, and also in the Lz of female placentas from malaria-infected dams. This pattern is consistent with findings in cytotrophoblast cultures isolated from human IUGR placentas (Vasilopoulou *et al*., 2010). MCT8 is expressed in both the syncytiotrophoblast and cytotrophoblast of the placental barrier, and its increased expression has been reported in IUGR placentas (Chan *et al*., 2006). An active compensatory mechanism of the maternal-fetal interface to enhance T3 uptake by trophoblast cells or facilitate the transplacental passage of iodothyronines has been previously suggested (Chan *et al*., 2006), demonstrating the importance of T3 metabolism and transport for appropriate fetal growth. Our data support the hypothesis of a compensatory mechanism, possibly stimulated by a local inflammatory process, typically observed in IUGR placentas with aberrant development of the fetoplacental unit resulting from reduced thyroid hormone supply by the trophoblasts (Loubière *et al*., 2010).

MCT10, initially characterized as a T-type amino acid transporter, is an active transporter of iodothyronines, with greater specificity for T3 when compared to MCT8 (Friesema *et al*., 2008). In the present study, we observed a downregulation in MCT10 protein expression in the placental labyrinth zone, as well as an upregulation in MCT10 protein expression in the placental junctional zone of both malaria-infected and LPS-injected dams. Unlike MCT8, the changes in MCT10 protein expression did not differ between sexes: expression was reduced in the Lz and increased in the Jz of both male and female offspring, in both experimental models. Since MCT10 is expressed in the syncytiotrophoblast and, mainly, in the cytotrophoblast of the placental labyrinth and its expression is also reduced in third-trimester IUGR placentas (Loubière *et al*., 2010), it is possible that the changes observed in MCT10 expression in both experimental models of the present study are also contributing to a reduction in the supply of iodothyronines and amino acids to the fetal compartment.

OATP4A1 is primarily expressed in the syncytiotrophoblast of placental chorionic villi, playing a fundamental role in the transport of organic anions and thyroid hormones from the maternal to the fetal compartment (Peng *et al*., 2020). In the present study, an increase in OATP4A1 expression was observed in both placental labyrinth and junctional of male offspring from malaria-infected dams, as well as in the labyrinth of both male and female offspring from LPS-injected dams. Given that the inhibition of OATP4A1 activity in human placentas reduces the transport of bile acids that are potentially detrimental to placental vascular development (Lofthouse *et al*., 2019), it is possible that the increased OATP4A1 expression observed in both experimental models of the present study could be associated with increased maternal-fetal transport of substrates potentially harmful to fetal development, thereby contributing to the observed IUGR.

The placental T3 accumulation also depends on the balance of D2 and D3 activities, the main iodothyronine deiodinases expressed in the human placenta (Chan *et al*., 2003). D2 is predominantly expressed in the cytotrophoblast, whereas D3 is mainly expressed in the syncytiotrophoblast (Chan *et al*., 2003). Interestingly, D2 and D3 expression are not altered in IUGR and preeclamptic human placentas (Chan *et al*., 2003; Kurlak *et al*., 2013). We did not observe changes in *Dio3* mRNA expression in either experimental model. In contrast, *Dio2* mRNA expression was reduced in the placentas of male offspring from malaria-infected dams.

D2 protein is regulated post-translationally by substrate-induced ubiquitination (Arrojo E Drigo & Bianco, 2011), this mechanism does no account for the reduction in Dio2mRNA observed here. Transcriptional repression by the local inflammatoty milieu is an alternative explanation that ramains to be tested, in special due to the reduced maternal fT4 observed in the MiP model. Interestingly, D2 may play a more significant role in the placenta than merely the conversion of T4 to T3. Downregulation of *Dio2* is observed in placentas of women with early recurrent miscarriages, possibly associated with reduced trophoblast invasive and migratory potential, leading to abnormal placentation (Adu-Gyamfi *et al*., 2021).

Overall, gestational inflammatory challenge was associated with a complex remodelling of the placental TH transport system. Both MiP and LPS increased MCT8 protein and produced a consistent zone-specific redistribution of MCT10, with reduced expression in the labyrinth and increased expression in the junctional zone, while LAT2 and OATP4A1 were increased in a sex-and compartment-dependent manner. At the transcriptional level, however, changes were limited, with reduced *Slc7a8* and *Dio2* expression confined to MiP. Together with the increased maternal TSH observed in both models and the reduced fT4 observed after MiP these findings indicate that gestational inflammation disrupts the maternal–placental TH axis through coordinated, sex-and compartment-dependent changes in TH transport and metabolism, although whether these alterations ultimately modify fetal TH availability remains to be determined.

The limitations of the present study include: the structural and functional differences between murine and human placentas (Dilworth & Sibley, 2013), which require caution regarding the translational potential of the results obtained in our experimental models; the administration of LPS, which, while well-established for mimicking gestational bacterial infections, elicits a purely endotoxic immune response (Reginatto *et al*., 2021), distinct from an infection with a pathogen, as seen in the MiP experimental model; and the inability to dissociate the effects of experimental cerebral malaria (ECM) from the onset of preterm labor starting at GD 17.5, even though the pregnant females were monitored daily between GD 13.5 and GD 18.5, confirming classic behaviors associated with ECM, such as ataxia and paralysis (De Oca *et al*., 2013).

In conclusion, this study demonstrates that murine models of MiP and acute endotoxemia induced by LPS generate distinct disturbances in the expression of placental TH transporters and deiodinases. Together with local inflammation and maternal thyroid dysfunction, the alterations may, to some extent, affect the placental regulation of iodothyronine transport and metabolism and contribute to adverse gestational outcomes such as fetal growth restriction and preterm delivery.

## CONFLICT OF INTEREST

There is no conflict of interest that could be perceived as prejudicing the impartiality of the research reported.

## AUTHOR CONTRIBUTIONS

**Klaus Novaes Fontes:** Conceptualization, Methodology, Formal Analysis, Investigation, Writing - Original Draft, Writing – Review & Editing, Project Administration. **Cherley Borba Vieira de Andrade:** Formal Analysis, Investigation, Writing – Review & Editing. **Verônica Muller de Oliveira Nascimento:** Formal Analysis, Investigation, Writing – Review & Editing. **Victoria Regina de Siqueira Monteiro:** Investigation, Writing – Review & Editing. **Hanailly Ribeiro Gomes:** Investigation, Writing – Review & Editing. **Alessandra Cristina Chagas Valim:** Investigation, Writing – Review & Editing. **Douglas Esteves Teixeira:** Methodology, Investigation, Writing – Review & Editing. **Ana Acácia Sá Pinheiro:** Methodology, Writing – Review & Editing, Resources. **Flavia Fonseca Bloise:** Resources, Writing – Review & Editing. **Enrrico Bloise:** Supervision, Writing – Review & Editing, **Tania Maria Ortiga-Carvalho:** Writing – Review & Editing, Resources, Project Administration, Funding Acquisition, Supervision

## DECLARATION OF GENERATIVE AI AND AI-ASSISTED TECHNOLOGIES IN THE MANUSCRIPT PREPARATION PROCESS

During the preparation of this work the author(s) used BioRender in order to create the chart of experimental design of both MiP and LPS models (Fig. 1). After using this tool/service, the authors reviewed and edited the content as needed and take full responsibility for the content of the published article.

## FUNDING

This work was supported by Conselho Nacional de Desenvolvimento Científico e Tecnológico (CNPq – grant n 309339/2023-5), Coordenação de Aperfeiçoamento Pessoal de Nível Superior (CAPES – Finance code 001) and Fundação de Amparo à Pesquisa do Estado do Rio de Janeiro (FAPERJ, grant nos: E26/201.206/2021; E26/201.957/2020; E26/2021.153/2022).

## ACKNOWLEDGEMENTS

We thank Juliana Gonçalves for her support with serum TH analysis assays, and Rakel Alves for her assistance with the animal experiments.

## ETHICS DECLARATION

This study was conducted in accordance with the ARRIVE (Animal Research: Reporting of In Vivo Experiments) guidelines as well as the following guidelines for animal welfare and/or reporting: “Principles of Laboratory Animal Care” - National Society for Medical Research and U.S. National Academy of Sciences Guide for the Care and Use of Laboratory Animals. This study was approved by the Animal Care Committee of the Health Sciences Center, Federal University of Rio de Janeiro.

## DATA AVAILABILITY

All data supporting the findings of this study are presented in the article. Raw data are available from the corresponding author on reasonable request.

## Notes

### Competing Interest Statement

The authors have declared no competing interest.

